# HY5 orchestrates the transcriptional network controlling coumarin-mediated iron acquisition under elevated pH conditions

**DOI:** 10.64898/2026.09.16.752116

**Authors:** Varsha Koolath, Muskan Kalra, Meijie Li, Samriti Mankotia, Anurag Kumar, Shunsuke Watanabe, Christian Dubos, Santosh B. Satbhai

**Affiliations:** Department of Biological Sciences, Indian Institute of Science Education and Research, Mohali, Punjab, 140306, India; IPSiM, Univ Montpellier, CNRS, INRAE, Institut Agro, Montpellier, France; Plant Science Research Laboratory (LRSV), CNRS, University of Toulouse, Toulouse INP, Toulouse, France; Department of Biological Science, Faculty of Science and Engineering, Yasuda Women’s University, Hiroshima, 731-0153 Japan

**Keywords:** HY5, coumarins, iron, pH, alkaline, transcriptional regulation, *Arabidopsis thaliana*

## Abstract

Iron (Fe) bioavailability is strongly limited in alkaline soils, where reduced Fe solubility severely restricts plant growth and productivity. To overcome this, *Arabidopsis thaliana* promotes Fe acquisition by secreting Fe-mobilizing coumarins, however, the transcriptional mechanisms coordinating this response is not clearly understood. Here we identify the ELONGATED HYPOCOTYL 5 (HY5), a bZIP transcription factor as a master regulator of coumarin mediated Fe acquisition under alkaline conditions. HY5 positively regulates genes required for coumarin biosynthesis, activation, secretion, and transcriptional control, and loss of HY5 markedly reduces their expression during high-pH-induced Fe deficiency. Chromatin immunoprecipitation analyses revealed that HY5 directly associates with the promoters of these genes, thereby establishing a transcriptional regulatory network that coordinates coumarin biosynthesis. Consistent with this regulatory function, *hy5* mutants exhibit reduced coumarin accumulation, impaired root growth, chlorosis, and decreased Fe accumulation under alkaline conditions. Comparable phenotypes in *coumarin-hy5* double mutants, together with the restoration of growth by exogenous fraxetin, demonstrate that HY5 functions upstream of coumarin-mediated Fe mobilization. Collectively, our findings identify HY5 as a molecular hub that integrates environmental pH signals with coumarin biosynthesis to promote adaptive Fe acquisition under alkaline conditions, providing a framework for improving crop performance on calcareous soils.

**SIGNIFICANCE STATEMENT:** Iron deficiency is a major limitation to crop productivity in alkaline soils, where iron is poorly soluble and difficult for plants to acquire. Although plants secrete iron-mobilizing coumarins to overcome this challenge, the regulatory mechanisms coordinating this adaptive response have remained largely unknown. We identify the transcription factor HY5 as a central regulator that directly coordinates coumarin biosynthesis, activation, secretion, and transcriptional control during iron deficiency at elevated pH. By linking environmental pH with coumarin-mediated iron acquisition, HY5 integrates external soil conditions with plant nutrient homeostasis. These findings reveal a previously unrecognized regulatory network underlying adaptation to alkaline soils and provide molecular targets for developing crops with improved iron-use efficiency and resilience in challenging agricultural environments.

## INTRODUCTION

As primary producers in the food chain, plants serve as a major source of non-heme iron for humans (1). With iron deficiency, humans are vulnerable to many health conditions, such as anaemia, which in severe cases, can even lead to organ failure (2, 3). In addition to anaemia, iron deficiency stands as a pervasive global health burden which comes with consequences that directly compromise the immune system of expectant mothers and children (4–6). It also drastically undermines physical productivity (7, 8) and inflicts lasting harm on the developing brains and nervous systems of affected individuals (9–11). Therefore, meeting the recommended daily iron intake in the human diet is essential. To ensure ample dietary iron availability, sufficient iron must accumulate in edible parts of crops, a goal that can be augmented by strategies such as biofortification (12).

Iron is also an essential micronutrient for plant growth and development (13). Iron homeostasis in plants involves iron uptake, transport and distribution, its use as a cofactor in metabolic processes, storage, and the tight molecular regulation of all four processes (14). Regardless of being the fourth most abundant element in Earth’s crust (15), iron uptake by plant roots is rather troublesome. That is because with each unit increase in pH between 4 and 9, the availability of iron decreases by 1000 fold (16, 17).

To overcome this hurdle, plants have evolved a range of strategies to efficiently mobilize rhizospheric iron. Non-graminaceous plants, such as *Arabidopsis thaliana*, employ a reduction-based mechanism to acquire iron from the rhizosphere under acidic conditions, called Strategy I. In such plants, the predominant form of iron on Earth’s crust, the ferric form (Fe^3+^), is reduced to available ferrous iron (Fe^2+^) by FERRIC REDUCTION OXIDASE 2 (FRO2) (18) after protonation of rhizosphere by AUTOINHIBITED PLASMA MEMBRANE H^+^ ATPase 2 (AHA2) (19), making Fe^2+^ available for uptake into roots through IRON REGULATED TRANSPORTER 1 (IRT1) (20).

In addition to the core reduction-based mechanism, Strategy I plants secrete iron-binding and mobilizing compounds into the rhizosphere when grown under iron-deficient conditions (21). Among these compounds are coumarins (22–26). In Arabidopsis, sideretin and fraxetin are the two main iron mobilizing coumarins (27) (28). Sideretin is mainly active at acidic pH and its secretion complements the classical Strategy I component FRO2 in reducing Fe³ in Fe² that is then up taken into the plant roots by IRT1 (27) (28). Under higher soil pH conditions, FRO2 activity becomes negligible, thereby limiting enzymatic Fe³ reduction (29). At this pH, it is proposed that fraxetin contributes to plant iron uptake in an IRT1-independent manner, by forming Fe^3+^-fraxetin complexes that are directly taken up by the plant roots through a mechanism that remains to be fully characterized (30).

Iron mobilising coumarins are biosynthesized from intermediates of the phenylpropanoid pathway through the sequential action of key enzymes, including FERULOYL-CoA 6′-HYDROXYLASE 1 (F6′H1), COUMARIN SYNTHASE (COSY), SCOPOLETIN 8-HYDROXYLASE (S8H), and the CYTOCHROME P450 MONOOXYGENASE CYP82C4 (31, 32). The pathway begins with F6′H1-mediated ortho-hydroxylation of feruloyl-CoA, followed by lactonisation, either spontaneously or catalysed by COSY, to yield the first stable coumarin, scopoletin (33, 34). S8H hydroxylates scopoletin to produce fraxetin (35–37), that is then converted into sideretin via CYP82C4 in acidic conditions (27). Once synthesized, these coumarins are stored in the vacuole as non-toxic glucosylated conjugates scopolin, fraxin, and sideretin glucoside (38–40). Before secretion into the rhizosphere, they undergo deglycosylation, after which the resulting aglycones are exported across the plasma membrane by the ATP-binding cassette (ABC) transporter PLEIOTROPIC DRUG RESISTANCE 9 (PDR9) (22, 41, 42).

Iron homeostasis is tightly controlled by a complex regulatory network in which bHLH transcription factors play a preponderant role. For instance, the expression of *FRO2* and *IRT1* is under the tight control of bHLH heterodimers composed of FIT/bHLH29 and clade Ib bHLHs, whose expression is under the control of URI/bHLH121 and clade IVc bHLHs (43–47). To prevent iron toxicity, this network is negatively regulated by HEMERYTHRIN E3-UBIQUITIN LIGASES such as BRUTUS (BTS). For instance, BTS function by targeting to the 26S proteasome degradation pathway URI/bHLH121 and two clade IVc bHLHs transcription factors, namely ILR3/bHLH105 and bHLH115. The negative regulator POPEYE/bHLH47 (PYE) also interact with ILR3 to inhibit the expression of genes involved in iron storage and transport, as well as its own expression (48–51). In addition, PYE also represses the expression of clade *Ib bHLHs* (52).

In contrast to the well elucidated transcriptional regulatory network that control the core Strategy I components, the regulation of coumarin biosynthesis and secretion remains only partially understood. For instance, it has been shown that the biosynthesis of iron mobilising coumarins requires URI, and that URI indirectly regulate the expression of *F6’H1*, *S8H* and *CYP82C4* (43, 53–56). MYB72, that is a direct target of URI, have been shown to play prominent roles in regulating the expression of coumarin-related genes under iron deficiency (28, 57). MYB63 was also shown to control the expression of *F6*′*H1*, *COSY* and *PDR9*, and therefore the secretion of coumarins, but under combined iron and phosphorus deficiency (58). While these transcription factors are known to contribute to coumarin-mediated iron mobilization, significant gaps remain in identifying master regulators that directly control the expression of genes involved in coumarin biosynthesis and secretion, and therefore that govern this key mechanism for plant iron nutrition.

The bZIP transcription factor ELONGATED HYPOCOTYL 5 (HY5) has recently emerged as a key positive regulator of iron acquisition in Arabidopsis, expanding its classical role as a master regulator of light-mediated photomorphogenesis and photomorphogenic development. We have previously reported that HY5 activates either indirectly or directly the expression of *FIT*, *FRO2*, and *IRT1* while repressing the expression of *BTS* and *PYE*, thereby fine-tuning iron-deficiency responses. In addition, we found reduced induction of coumarin-related genes in the *hy5* mutant compared to wild-type plants under iron-deficient conditions, raising the intriguing possibility that HY5 contributes to coumarin biosynthesis regulation (59, 60).

Through a comprehensive integration of molecular, genetic, physiological, and biochemical assays, we investigated HY5’s contribution to coumarin-mediated iron uptake. Spatial colocalization of HY5 and target gene reporters, together with reduced coumarin accumulation and secretion in *hy5* plants, confirms that HY5 is required for optimal coumarin production. Genetic analyses combined with physiological assays under varying pH and iron availability, establish that the HY5-coumarin module is essential for maintaining root growth and iron homeostasis, particularly under neutral to alkaline pH, when the enzymatic reduction of Fe³ by FRO2 is markedly impaired (61). ChIP-qPCR confirms that HY5 directly binds to and positively regulates the expression of all the key coumarin pathway genes, including *PDR9* and *MYB72*. Collectively, these findings position HY5 as a central integrator of iron acquisition, revealing a molecular link between environmental cues and the mobilization of rhizosphere nutrients under adverse conditions. This regulatory mechanism offers promising avenues for enhancing the efficiency of iron homeostasis and biofortification in crops grown on calcareous, high-pH soils, where iron deficiency severely constrains productivity.

## RESULTS

### Disruption of HY5 impairs the expression of genes involved in iron-mobilizing coumarins pathway

To investigate the role of *HY5* in coumarin biosynthesis, we analyzed the expression of key iron-mobilizing coumarin biosynthetic genes in wild-type (WT) and *hy5* mutant seedlings. It has been reported that the expression of coumarin pathway genes was less induced in *hy5* under iron deficiency at acidic pH (pH 5.7) (59). RT-qPCR results confirmed previous observation and revealed reduced induction of *F6’H1*, *S8H* and *CYP82C4* expression in the *hy5* mutant compared to WT plants at high pH (pH 7) (Fig. 1*A*). To determine whether this regulatory effect extended beyond the core biosynthetic pathway, we also checked the expression of genes involved in coumarin secretion, *BGLU42* and *PDR9* (*SI Appendix*, Fig. S1) and transcriptional regulation, *MYB72*. Since *MYB10* was shown to act redundantly with *MYB72* in promoting plant growth under alkaline condition, its expression was also studied (*SI Appendix*, Fig. S2). The reduced expression of *BGLU42*, *PDR9*, *MYB10* and *MYB72* in the *hy5* mutant indicates a broader role for HY5 in coordinating the transcriptional network underlying coumarin-mediated Fe mobilization, particularly under elevated pH conditions.

**Figure 1.**
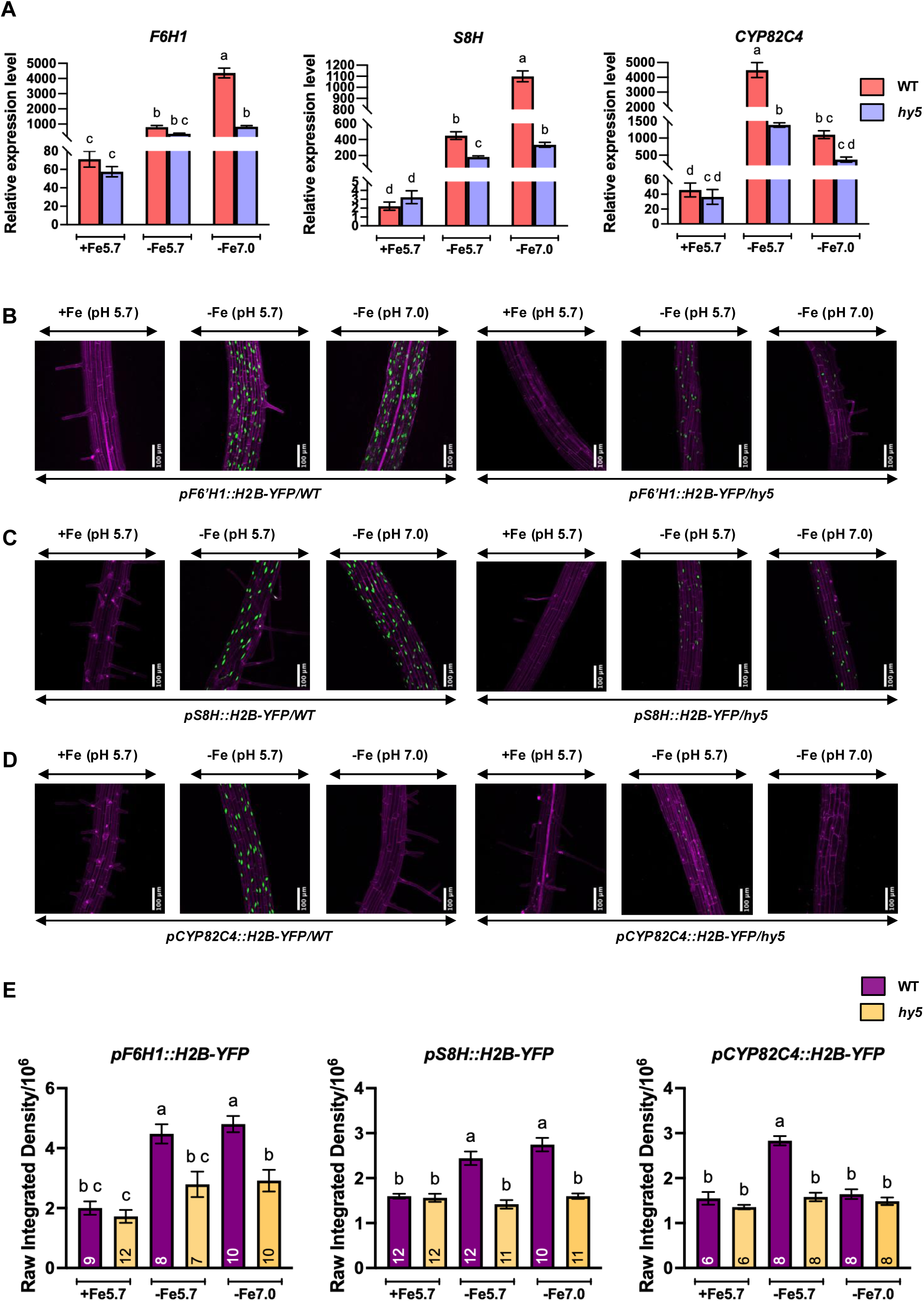
*HY5* positively regulates the expression of coumarin biosynthesis genes. (*A*) Gene expression profiles (RT qPCR) of coumarin biosynthesis genes *F6’H1*, *S8H,* and *CYP82C4* in the roots of Col-0 and *hy5* mutant seedlings grown on +Fe pH 5.7 media for 6 days and then transferred to +Fe pH 5.7, -Fe pH 5.7, and -Fe pH 7.0 for 3 days. Transcript levels were normalized to *PP2AA3*. Data are mean ± SEM of three biological replicates (two technical replicates each); each biological replicate is pooled RNA from ∼100 roots. Different letters indicate significant differences (one-way ANOVA, Tukey’s post hoc test, p < 0.05, GraphPad Prism 11). (*B–D*) Confocal images of *F6’H1::H2B-YFP* (*B*), *S8H::H2B-YFP* (*C*), and *CYP82C4::H2B-YFP* (*D*) reporter signal in WT and *hy5* backgrounds. Five days old seedlings grown on +Fe (pH 5.7) medium were transferred onto +Fe (pH 5.7) and -Fe (pH 5.7 and pH 7.0) media for three days, and then stained with 10 µM propidium iodide prior to imaging of the maturation zone. Green: YFP, magenta: propidium iodide. Confocal imaging was performed in Z-stack mode with a 1 µm step size. Scale bar = 100 µm. (*E*) Quantification of raw integrated density of YFP signals in *F6’H1::H2B-YFP*, *S8H::H2B-YFP,* and *CYP82C4::H2B-YFP* in WT and *hy5* mutant background. Error bars represent ±SEM. n is annotated at the bottom of each bar. Different letters indicate a significant difference according to one-way ANOVA followed by post hoc Tukey test, p < 0.05, using GraphPad PRISM 11.

To confirm these observations *in planta*, we generated transcriptional H2B-YFP reporter lines driven by the native promoter of *F6’H1*, *S8H* and *CYP82C4* in Col-0 and *hy5* mutant backgrounds. As expected, YFP signals were detected in the nuclei of cells, predominantly in the maturation zone (28). In the *hy5* mutant background, the YFP signal was reduced for all three coumarin biosynthetic genes under iron-limited conditions at both pH 5.7 and pH 7.0 (Fig. 1*B-E*).

Taken together, these findings confirm that HY5 positively regulates the expression of the key coumarin biosynthesis genes *F6’H1*, *S8H,* and *CYP82C4*, and thereby enhances the transcriptional activation of the Fe-mobilizing coumarin pathway under alkaline conditions. It also supports that HY5 regulate the expression of genes involved in coumarin secretion and the regulation of these processes.

### HY5 directly regulates the expression of genes involved in iron-mobilizing coumarins pathway

The reduced expression of coumarin biosynthetic genes in the *hy5* mutant indicated that HY5 may directly regulate their transcription. To confirm this hypothesis, we first performed *in silico* analysis of the promoter sequences of *F6’H1*, *S8H,* and *CYP82C4* for the presence of known HY5 binding motifs (Fig. 2*A*). Multiple canonical HY5 binding motifs, including G-box (*CACGTG*), specific E-box (*CAATTG*) and CA-Hybrid element (*GACGTA*), were identified within the promoters of all three genes (62–66). Specifically, the promoter of *F6’H1* contained three DNA binding motifs (one E-box and two G-boxes), the promoter of *S8H*, four (two CA-Hybrid, one E-box and one G-box), and the promoter of *CYP82C4*, only one (CA-Hybrid motif) (Fig. 2*A*). To determine whether HY5 directly associates with these promoter regions *in vivo*, we performed chromatin immunoprecipitation followed by quantitative PCR (ChIP-qPCR). HY5 was significantly enriched at the promoters of *F6’H1*, *S8H*, and *CYP82C4*, demonstrating direct binding to these regulatory regions (Fig. 2*B–I*). These findings establish that HY5 directly activates the transcription of these key iron-mobilizing coumarin biosynthesis genes.

**Figure 2.**
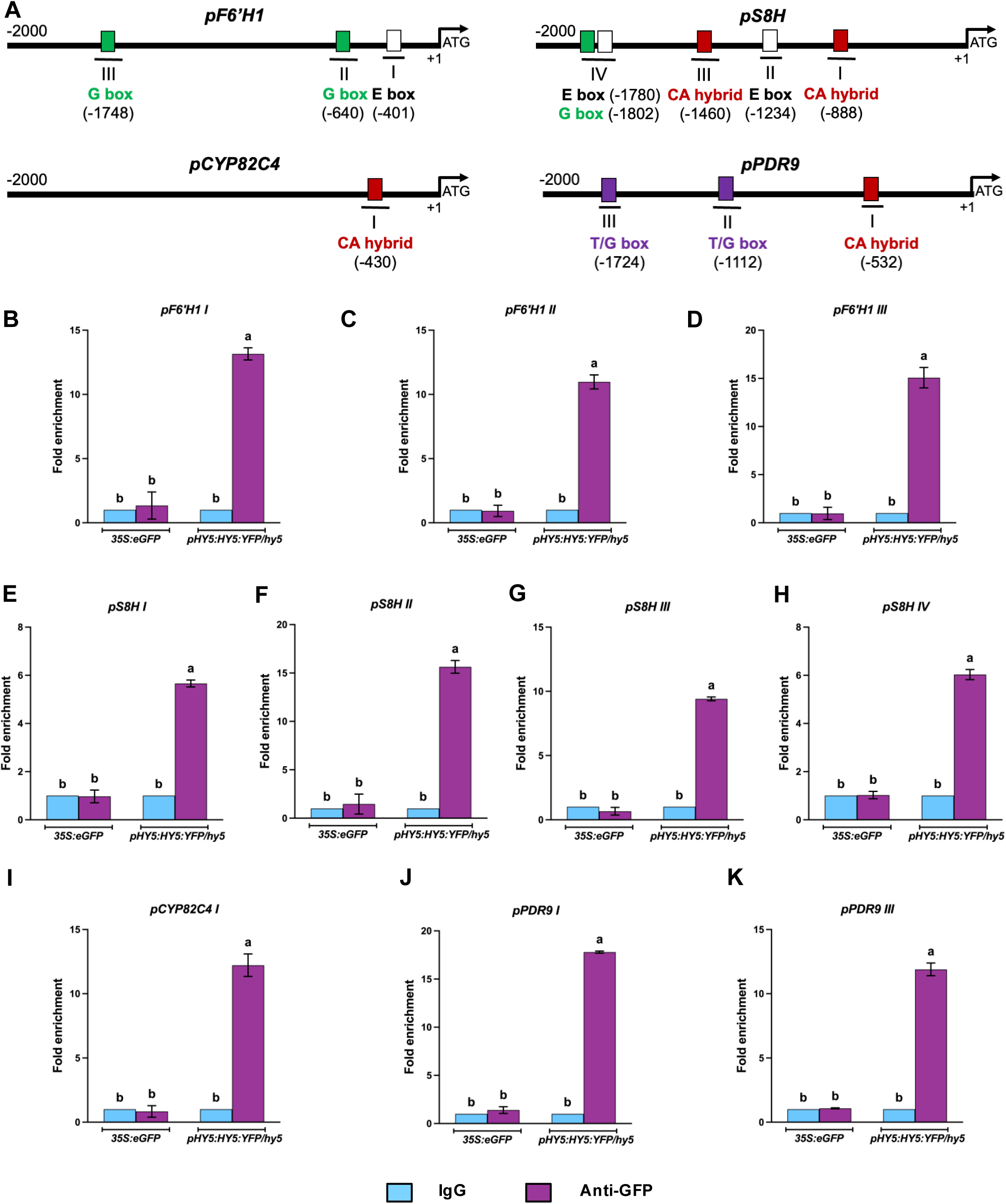
HY5 directly binds on promoters of coumarin biosynthesis and secretory genes *F6’H1*, *S8H*, *CYP82C4* and *PDR9*. *A*) Schematic representation of the *F6’H1*, *S8H*, *CYP82C4* and *PDR9* promoter regions. Lines beneath the boxes denote the regions analyzed by ChIP-qPCR. The numbers in bracket denotes the number of bases upstream to the respective gene’s start codon. *B-K*) ChIP-qPCR analysis showing the enrichment of HY5 at the binding motifs of *F6’H1*, *S8H*, *CYP82C4* and *PDR9* promoter. The ChIP assay was performed using *pHY5::HY5:YFP/hy5* (Ler) and *35S::eGFP* (Ler) seedlings grown on +Fe medium for 10 days. n=2 biological replicates for each genotype. Error bars represent ±SEM. Different letters indicate significant difference according to one-way ANOVA followed by post hoc Tukey test, p< 0.05. using GraphPad PRISM 11.

Because *BGLU42*, *PDR9*, *MYB10,* and *MYB72* also exhibited reduced expression in the *hy5* mutant, we next examined whether these genes are also direct HY5 targets. Promoter sequence analysis identified multiple potential HY5-binding motifs, including E-box, G-box, and CA-hybrid box, as well as T/G-box (*AACGTG*), A-box (*TACGTA*), Z-box (*ATACGTGT*), and GATA-box (*GATGATA*) (Fig. 2*A*; *SI Appendix*, Fig. S1 and Fig. S2) (62–66). Consistent with our predictions, ChIP-qPCR analysis revealed significant enrichment of HY5 at the promoters of *PDR9* (Fig. 2*J–K*) and BGLU42 (*SI Appendix*, Fig. S1) as well as *MYB10* and *MYB72* (*SI Appendix*, Fig. S2), indicating direct promoter occupancy by HY5.

In summary, these results demonstrate that HY5 directly binds to and regulates both structural (*F6’H1*, *S8H*, *CYP82C4*, *BGLU42* and *PDR9*) and regulatory (*MYB10* and *MYB72*) components of the coumarin pathway, thereby positioning HY5 as a central transcriptional regulator of coumarin-mediated iron acquisition.

### HY5 is required for iron-mobilizing coumarins biosynthesis and secretion

To determine whether the transcriptional regulation mediated by *HY5* influences coumarin production and secretion, we examined coumarin accumulation in WT and *hy5* seedlings grown under iron-limiting conditions. Coumarin biosynthesis and secretion was visualized using UV fluorescence and biphoton confocal microscopy, followed by linear spectral unmixing. Additionally, phenolic levels were quantified via high-performance liquid chromatography (HPLC). When compared to the root of WT plants grown at pH 5.7, the roots of plants grown at high pH (7.0) display prominent UV-fluorescent signals corresponding to the coumarins present in the rhizosphere. This observation is consistent with the role of fraxetin in chelating and mobilizing insoluble Fe^3+^. In contrast, in the *hy5* mutant, fluorescence intensity was highly reduced (Fig. 3*A*). Consistently, biphoton confocal imaging revealed substantially lower fraxin accumulation in *hy5* roots than in WT at pH 7, as well as lower accumulation of its precursor scopolin, at both pH (Fig. 3*B*).

**Figure 3.**
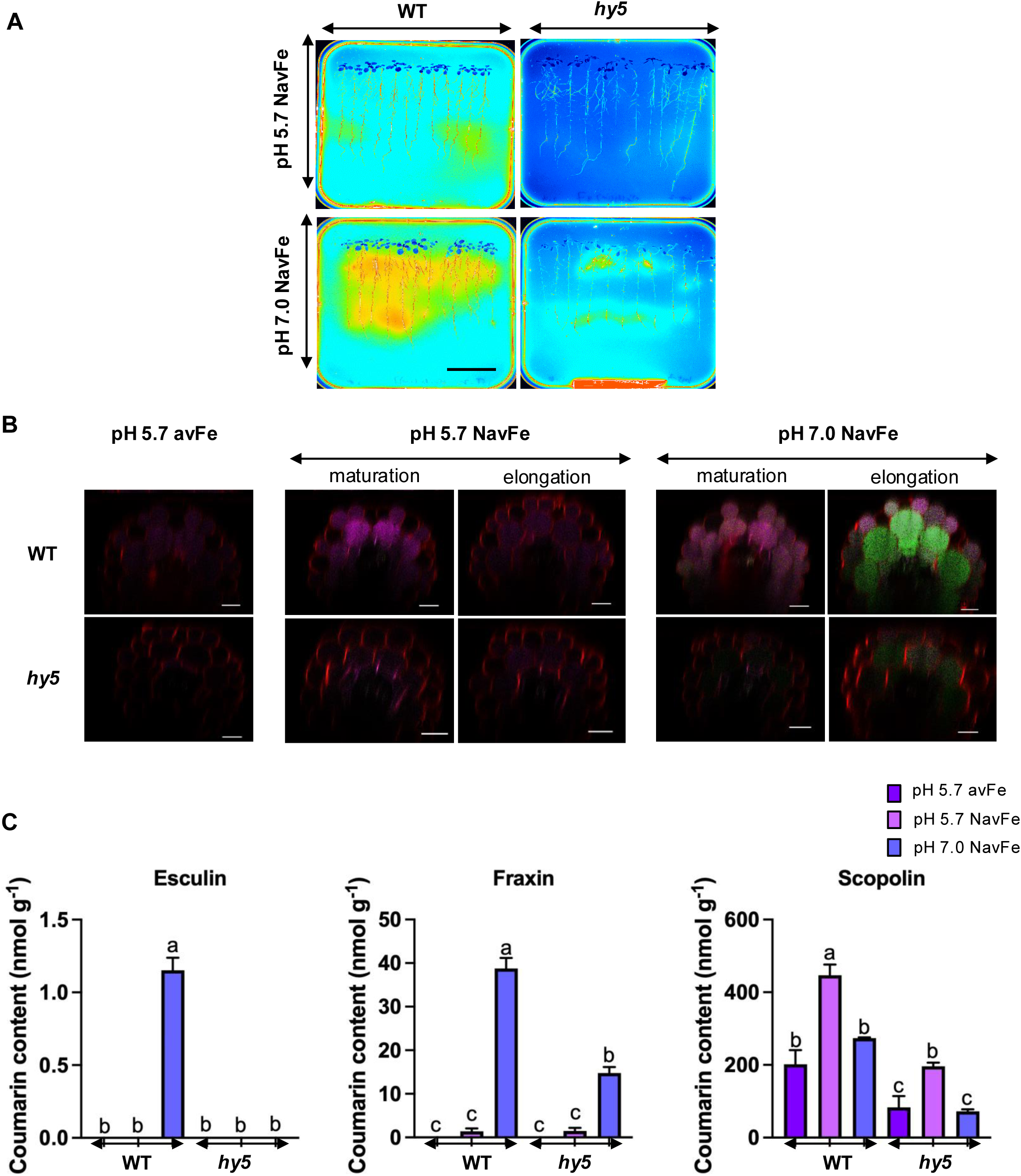
*HY5* is required for coumarin secretion and accumulation at elevated pH. (*A*) Coumarin secretion (UV autofluorescence, 365 nm) of wild type (WT) *and hy5* mutants grown in control condition (50 µM Fe-EDTA, pH 5.7) for 9 days and then transferred to media that contain a form of iron that is poorly available to plants (i.e., 100 µM FeCl_3_), at pH 5.7 (NavFe pH 5.7) or pH 7 (NavFe pH 7.0), for 3 additional days. (*B*) Coumarin distribution and localization in the *hy5* mutant. Images were obtained by linear unmixing with the emission spectra of scopolin (purple), fraxin (green), esculin (dark yellow), and propidium iodide (PI, red). Seedlings were grown in control conditions (i.e., 50 µM Fe-EDTA, pH 5.7) for 5 days and then transferred to control or media containing poorly available iron (i.e., 100 µM FeCl_3_), at pH 5.7 or pH 7, for 6 additional days. Scale bar=20 µm. (*C*) Esculin, scopolin and fraxin contents in WT and *hy5* seedlings. Means within each condition with the same letter are not significantly different according to one-way ANOVA followed by post hoc Tukey test, *p* < 0.05 using GraphPad Prism11.

HPLC analysis further confirmed a significant reduction in accumulation of the major coumarin glucosides, including esculin, fraxin, and scopolin in the *hy5* mutant relative to WT (Fig. 3*C*). The significantly reduced coumarin levels were consistent with the reduced expression of coumarin biosynthetic genes observed by RT-qPCR, indicating that the loss of *HY5* severely affects both coumarin biosynthesis and secretion. Collectively, these findings demonstrate that *HY5* is required for the efficient production and secretion of Fe-mobilizing coumarins, particularly under alkaline conditions, supporting its vital role in the coumarin-dependent iron acquisition pathway.

### Genetic studies confirm that HY5 acts upstream from genes involved in iron-mobilizing coumarins pathway

To elucidate the genetic relationship between *HY5* and coumarin pathway genes, we generated double mutants by crossing *hy5* with T-DNA insertion mutants of *F6’H1, S8H, CYP82C4, PDR9*, and *MYB72*. Homozygous double mutants were confirmed by genotyping and subsequently characterized for primary root growth, chlorophyll accumulation, and Fe content under different Fe availability and pH conditions. No pronounced growth differences were observed at pH 5.7 between *hy5* and all double mutants (*f6’h1 hy5*, *s8h hy5*, *cyp82c4 hy5*, *pdr9 hy5*, and *myb72 hy5*) (Fig. 4*A-E*; *SI Appendix*, Fig. S4). At pH 7, *hy5* and all double mutants exhibited significantly shorter primary roots than WT plants. In contrast, chlorophyll content was reduced in double mutants, reflecting potential photosynthetic defects associated with iron deficiency (*SI Appendix*, Fig. S3 and Fig S4). Perls’ staining revealed reduced blue precipitates in root tissues of both *hy5* single mutant and all double mutants, signifying decreased iron accumulation. At pH 5.7, WT and coumarin single mutants showed more iron accumulation compared to the *hy5* single mutant and all the double mutants. At pH 7.0, we observed the least staining in *hy5* single mutants and all double mutants, followed by coumarin single mutants (*f6’h1* and *s8h*), and the highest in *cyp82c4* and WT (*SI Appendix*, Fig. S5). Comparable staining patterns were observed for both Perls’ and Perls’-DAB assays. Similar phenotypes were also observed in the *pdr9 hy5* and *myb72 hy5* double mutants and the total iron content estimation outcome was consistent with Perls’ and Perls’ DAB staining results (*SI Appendix*, Fig. S5 and Fig. S6).

**Figure 4.**
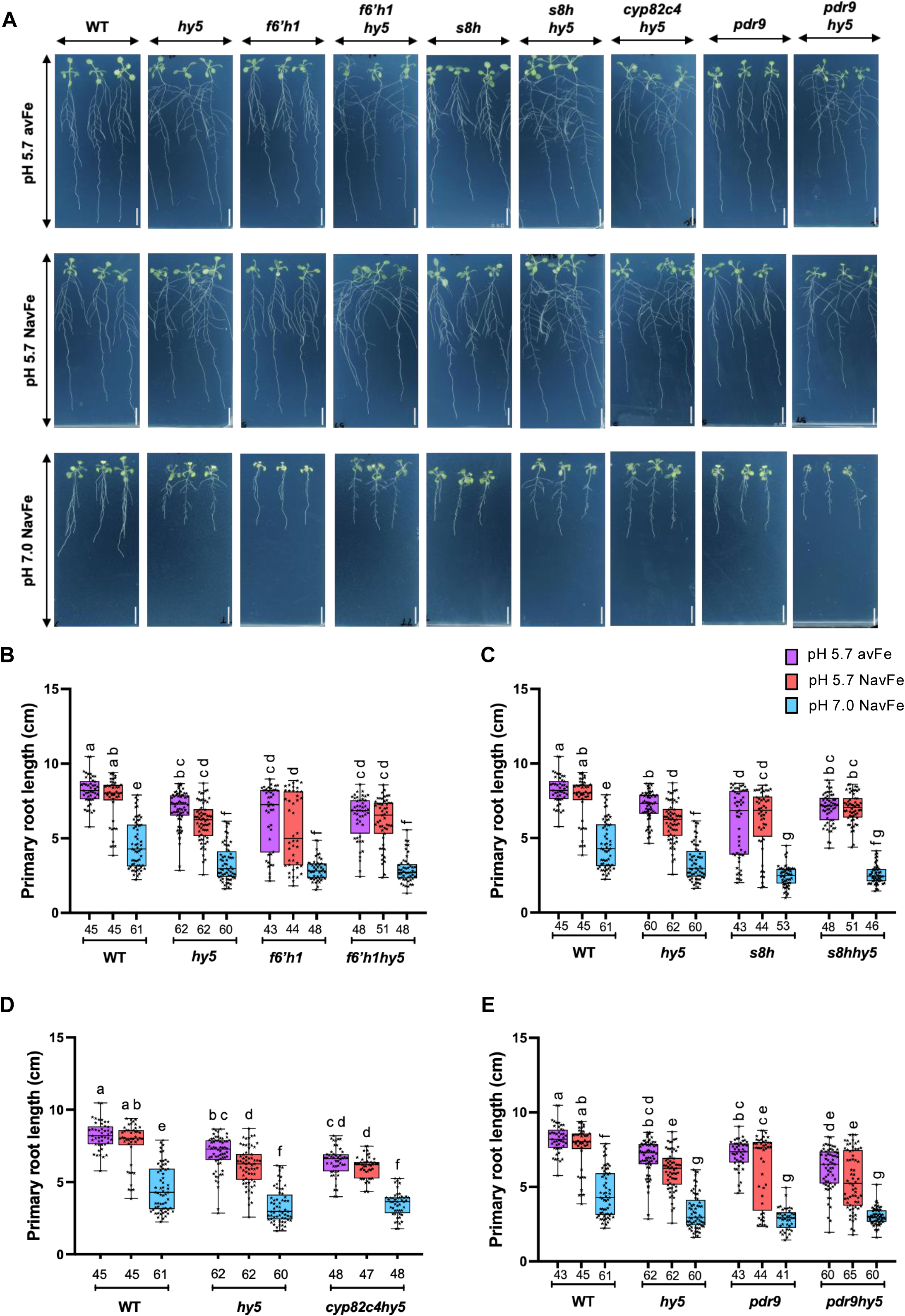
*HY5* genetically interacts with coumarin biosynthesis and secretion genes at high pH. *A*) Phenotype of WT*, hy5, f6’h1, f6’h1 hy5, s8h*, *s8h hy5*, *cyp82c4 hy5, pdr9* and *pdr9 hy5* seedlings grown in contrasted iron nutrition. Five days old seedlings grown at pH 5.7 on avFe were transferred to avFe at pH 5.7 and NavFe media at both pH 5.7 and pH 7.0 and grown further for nine days before data acquisition. Scale bar=1cm. (*B-D*) Primary root length of coumarin biosynthesis-gene *hy5* double mutants: *f6’h1 hy5* (*B*), *s8h hy5* (*C*), *cyp82c4 hy5*(*D*), under the conditions in (*A*). n indicated on the x-axis.(*E*) Primary root length of the coumarin secretion-gene double mutant *pdr9 hy5*, under the conditions in (*A*). n indicated on the x-axis. Means with the different letters are significantly different according to ordinary two-way ANOVA followed by post-hoc Tukey test, p < 0.05, using GraphPad PRISM 11.

Altogether, these genetic analyses demonstrate that disruption of coumarin biosynthesis, secretion, or transcriptional regulation enhances the Fe-deficiency phenotypes associated with *hy5* under elevated pH conditions. The observed epistatic relationships support a model in which HY5 functions upstream of the iron-mobilizing coumarin pathway to promote iron acquisition during alkaline stress.

### HY5 and proteins involved in iron-mobilizing coumarins pathway accumulate in the same root cell types

To examine the spatial relationship between HY5 and the coumarin pathway, we analyzed F1 progeny generated by crossing stable transgenic reporter lines expressing native promoter-driven HY5-YFP with reporter lines carrying the whole *F6’H1*, *S8H*, *CYP82C4*, and *PDR9* genomic loci fused to fluorescent (RFP or mCitrine) reporters. Confocal imaging showed that nuclear-localized HY5-YFP signals overlapped with the localization of all five coumarin gene reporter signals in the same root cell layers (Fig. 5*A-D*). Co-expression was predominantly observed in the epidermis and cortex of elongation and maturation zones. The localization pattern aligns with known iron-acquisition zones in the root, where outer cell layers coordinate iron sensing, mobilization, and uptake (30, 31, 33). We also checked the localization pattern of BGLU42 and HY5 (*SI Appendix*, Fig. S7). Notably, the overlap in protein localization was enhanced under alkaline iron-limiting conditions consistent with the induction of the coumarin pathway during Fe deficiency, except for CYP82C4 whose expression is repressed at this pH (67, 68). The spatial co-localization of HY5 with coumarin biosynthesis and secretion genes, together with the reduced expression of these genes in the *hy5* mutant, strongly supports a role for HY5 as a local transcriptional regulator coordinating coumarin production in root tissues responsible for iron mobilization, particularly under alkaline conditions.

**Figure 5.**
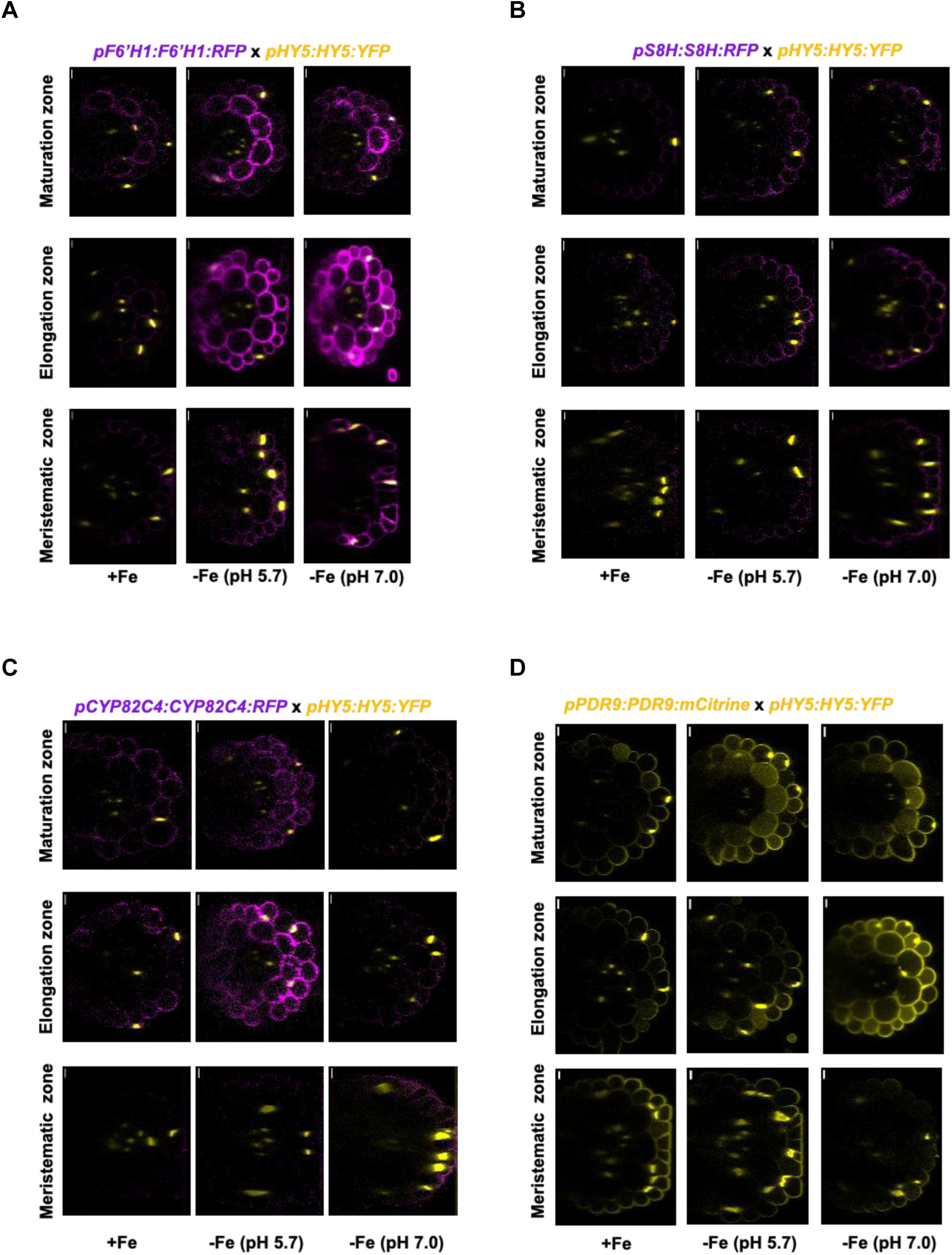
HY5 expression colocalizes with the expression of the genes involved in the coumarin biosynthesis and secretion. *A-D*) The seedlings of HY5 translational YFP fusion line crossed with F6’H1 (*A*), S8H (*B*), CYP82C4 (*C*) translational RFP fusion lines and PDR9 translational mCitrine fusion lines (*D*) were grown on +Fe medium for four days and transferred to -Fe medium at pH 5.7 and pH 7.0 for additional 3 days. Yellow: YFP or mCitrine fluorescence, magenta RFP. Confocal imaging was performed in Z-stack mode with a step size of 1 µm. Scale bar= 10 µm.

### Fraxetin-mediated rescue places HY5 upstream of coumarin-mediated iron acquisition

In order to investigate whether phenotypes of *hy5* mutants under high pH and Fe-limiting conditions are primarily due to impaired coumarin production, seedlings were grown at pH 7 in the presence of exogenous fraxetin. Supplementation of exogenous fraxetin significantly restored primary root growth and alleviated leaf chlorosis in *hy5*, coumarin single mutants, and the corresponding double mutants, except for *cyp82c4* that has enhanced fraxetin biosynthesis (Fig. 6*A-B*; *SI Appendix*, Fig. S8). In contrast, solvent controls had no detectable effect, confirming that the observed rescue was specific to fraxetin treatment. The restoration of root growth and chlorophyll levels by exogenous fraxetin indicates that the developmental and physiological defects of the *hy5* mutant under alkaline iron limitation mainly arise from impaired endogenous coumarin biosynthesis. In other words, this finding shows that externally supplied fraxetin bypasses the requirement for HY5-dependent activation of the coumarin pathway, thereby restoring efficient iron acquisition under high-pH conditions.

**Figure 6.**
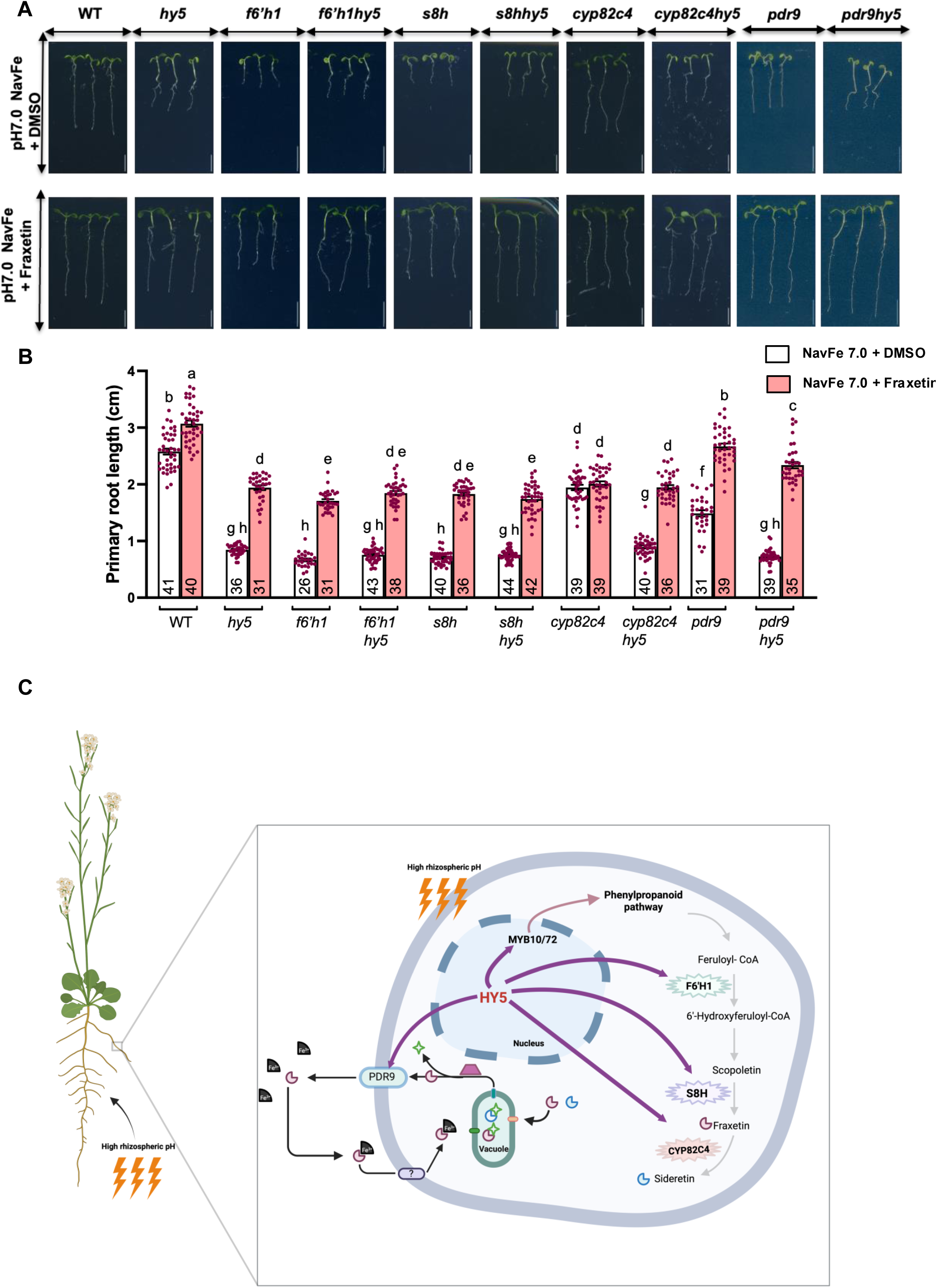
Fraxetin supplementation restores root growth under high pH mediated iron deficiency. *A*) Phenotype of WT*, hy5, f6’h1, f6’h1 hy5, s8h, s8h hy5, cyp82c4, cyp82c4 hy5*, *pdr9* and *pdr9 hy5* seedlings grown on NavFe media at pH 7.0. for four days and transferred to NavFe at pH 7.0 media with fraxetin or solvent control (DMSO) and grown further for three days before data acquisition. Scale bar=1cm. *B*) Primary root length of plants described in (*A*). Means with the different letters are significantly different according to one-way ANOVA followed by post-hoc Tukey test, p < 0.05, using GraphPad PRISM 11. n is annotated within the bar at the bottom. Error bars represent ±SEM. *C*) Model: Under high rhizospheric pH conditions, the bZIP transcription factor HY5 acts as a central regulator orchestrating both coumarin biosynthesis and iron uptake responses in *Arabidopsis thaliana* root cells. HY5, operating within the nucleus, acts in concert with MYB10/72 to modulate the iron deficiency response. HY5 promotes the phenylpropanoid pathway, leading to the sequential conversion of feruloyl-CoA to 6’-hydroxyferuloyl-CoA via F6’H1, followed by the production of scopoletin, which is further hydroxylated by S8H to yield fraxetin under high rhizospheric pH conditions, and subsequently converted to sideretin by CYP82C4 under lower rhizospheric pH by direct promoter occupancy interactions. Coumarins are stored in vacuoles as coumarin-glucoside until deglycosylation for secretion in response to iron deficiency. HY5 directly activates the expression of *PDR9*, encoding the efflux transporter responsible for secretion of coumarins into the rhizosphere. Once secreted, coumarins chelate and help mobilize sparingly soluble Fe^3+^ in the rhizosphere, facilitating its subsequent reduction and uptake into root cells as either Coumarin-Fe^3+^ conjugates or Fe^2+^. The model was adapted from (68) and created with BioRender.

## DISCUSSION

Iron deficiency is a major constraint on plant productivity in alkaline and calcareous soils, where ferric iron is predominantly present as insoluble oxyhydroxides. Under these conditions, the canonical Strategy I iron uptake system becomes progressively less effective because ferric reduction is strongly inhibited by elevated pH (18, 31). Arabidopsis and other dicot species overcome this limitation by secreting iron mobilizing coumarins, specialized phenolic metabolites that chelate and reduce ferric iron, thereby increasing its bioavailability in the rhizosphere for root uptake (13, 31, 69). Although the enzymatic steps underlying coumarin biosynthesis have been extensively characterized (27, 33, 35, 36), the transcriptional mechanisms that coordinate this pathway in response to alkaline conditions have remained poorly understood (28, 44, 67, 68). Here, we identify HY5 as a central transcriptional regulator that coordinates coumarin biosynthesis and secretion, as well as regulatory networks, to enable iron acquisition under alkaline conditions.

Our data demonstrate that HY5 directly activates the expression of all known major components of the coumarin pathway required for iron mobilization. *HY5* positively regulates the expression of the core biosynthetic genes *F6*′*H1*, *S8H*, and *CYP82C4* (Fig. 1*A-E*, Fig. 2*B-I*), the secretion-associated genes *BGLU42* and *PDR9* (Fig. 2*J-K*; *SI Appendix*, Fig. S1), and the transcriptional regulators *MYB72* and *MYB10* (*SI Appendix*, Fig. S2). Notably, direct HY5 occupancy at the *PDR9* promoter identifies the terminal transport step, rather than only the upstream biosynthetic and regulatory nodes, as a *bona fide* component of this transcriptional circuit, indicating that HY5 governs coumarin-dependent iron mobilization at the point of rhizosphere delivery (Fig. 2*J-K*). This direct control extends beyond structural pathway genes to the regulatory tier itself: HY5 also directly targets MYB72, suggesting that HY5 reinforces coumarin production through a coherent feed-forward transcriptional circuit in which activation of a secondary transcription factor amplifies downstream metabolic output (*SI Appendix*, Fig. S2). Thus, rather than controlling a single biosynthetic step, HY5 coordinates the complete biosynthetic, regulatory, and secretion machinery required for coumarin-dependent iron mobilization.

This regulatory function of HY5 is reflected at the metabolic and physiological levels. Independent fluorescence imaging approaches, together with HPLC-based metabolite profiling, revealed a marked reduction in coumarin accumulation in *hy5* mutants under alkaline conditions, demonstrating that HY5-dependent transcription translates into metabolic flux through the coumarin pathway (Fig. 3*A-C*). Consistent with these observations, *hy5* double mutants with gene from the coumarins pathway displayed phenotypes comparable to the *hy5* single mutant with respect to root growth, chlorophyll accumulation, and iron content under high pH iron-limiting conditions, supporting a linear regulatory hierarchy in which *HY5* acts upstream of the coumarin pathway (Fig. 4*A-E*, *SI Appendix*, Fig. S3, S4, S5, and S6). These genetic relationships were most pronounced under iron-limiting alkaline conditions, emphasizing that HY5 becomes particularly important when conventional ferric reduction is compromised by elevated pH.

Our spatial expression analyses further strengthen this model. HY5 and its downstream targets were co-expressed predominantly in the epidermis and cortex of the elongation and maturation zones, the main sites of coumarin synthesis and secretion during iron deficiency, particularly under high-pH conditions. Moreover, the expression of both HY5 and its targets was substantially enhanced at pH 7.0 compared with pH 5.7, indicating that activation of this transcriptional module is activated in response to changes in external pH (Fig. 5*A-D*; *SI Appendix*, Fig. S7). These observations place HY5 at the interface between environmental sensing and localized metabolic responses that promote rhizosphere iron mobilization.

The physiological importance of this pathway is further shown by the restoration of growth and chlorophyll accumulation following exogenous fraxetin application. Fraxetin rescued the growth and chlorophyll defects of *hy5*, coumarin-pathway mutants, and their corresponding double mutants under alkaline iron deficiency, indicating that the defects associated with *HY5* loss largely result from impaired coumarin production. Because externally provided fraxetin bypasses the requirement for endogenous biosynthesis, these findings further support that *HY5* acts upstream of coumarin-mediated iron mobilization (Fig. 6*A-B*; *SI Appendix*, Fig. S8). These findings also reinforce the central role of coumarins in facilitating iron acquisition under conditions in which the classical ferric-reduction pathway is ineffective.

Collectively, our results support a model in which alkaline pH activates a *HY5*-dependent transcriptional program that co-ordinately induces coumarin biosynthesis (*F6*′*H1*, *S8H*, *CYP82C4*), coumarin activation (*BGLU42*), secretion (*PDR9*), and pathway amplification through *MYB72* and *MYB10*. The resulting accumulation and export of fraxetin and related coumarins enhance the mobilization of insoluble ferric iron in the rhizosphere, thereby complementing the canonical FIT–IRT1/FRO2 iron uptake system that predominates under mildly acidic conditions. These findings substantially expand the known functions of HY5 by identifying it as a molecular hub that integrates environmental pH with adaptive metabolic responses required for iron acquisition (Fig. 6*C*).

Several important questions remain. How alkaline pH activates HY5, whether HY5 functionally interacts with the URI, FIT and other bHLH from the iron-deficiency regulatory network under alkaline conditions, whether this HY5-dependent module extends to coumarin-dependent induced systemic resistance, a process that shares several of the same regulatory and secretion components, and how this regulatory circuit operates in heterogeneous field soils remain to be determined. It will also be important to establish whether analogous transcriptional mechanisms regulate phenolic-mediated iron acquisition in other Strategy I species or in grass crops that uses the Strategy II (chelation strategy). From a translational perspective, the HY5-centered regulatory module provides an attractive target for engineering crops with enhanced tolerance to alkaline soils. Manipulating HY5 activity or redesigning HY5-responsive promoters could co-ordinately activate the entire coumarin biosynthetic and secretion network, offering a systems-level approach to improving iron acquisition and nutritional resilience in agricultural environments with increasingly widespread soil alkalization.

## MATERIALS AND METHODS

### Plant materials and growth conditions

In this study, the Columbia (Col-0) ecotype of *Arabidopsis thaliana* was used as the WT. The mutant/transgenic lines used in this study were *hy5* [SALK_056405; (70)], *f6’h1-1* [At3g13610, SALK_132418C; (33)], *s8h-2* [At3g12900, SM_3.23443; (36)], *cyp82C4* [At4g31940, SALK_001585; (27)], *pdr9-2* [At3g53280, SALK_050885; (42)], *myb72* [At1g56160, SAIL_713_G10C, CS867464] and *pPDR9:PDR9:mCitrine* in *pdr9-2* (31). *pHY5:HY5:YFP/hy5* (Ler) and *35S:eGFP* (Ler) seeds were provided by Roman Ulm. Seedlings were grown under long day conditions, 16 h light and 8 h dark at 22°C with a light intensity of 100-110 μmol cm^-2^ s^-1^ and 50% humidity. The soilrite, perlite and compost (3:1:0.5) mixture was used as soil. To generate the double-mutants, *f6’h1 hy5*, *s8h hy5*, *cyp82C4 hy5*, *pdr9 hy5* and *myb72 hy5*, *hy5* single-mutant was crossed with *f6’h1*, *s8h*, *cyp82C4*, *pdr9, myb72* single mutants respectively. The single as well as double mutants were confirmed by PCR genotyping. The primers used for genotyping are listed in *SI Appendix*, Table S1.

### Gene Expression analysis

Total RNA was extracted from seedlings grown on 0.5 MS medium for 6 days and then transferred to iron-sufficient (+Fe 5.7), iron-deficient (–Fe 5.7 and –Fe 7.0) for 3 days using the NucleoSpin RNA extraction kit (Macherey-Nagel). For each sample, 1 µg of total RNA treated with DNase was used to synthesize cDNA with the RevertAid kit (Thermo Scientific). qRT PCR analyses were performed using a LightCycler 480 (Roche) and TB Green Premix Ex Taq (2X; Takara). *PP2AA3* (PROTEIN PHOSPHATASE 2A SUBUNIT A3) was used as a reference gene (71). Expression levels were calculated using the comparative threshold cycle method. The primers used for genotyping are listed in *SI Appendix*, Table S1.

### Cloning

To generate coumarins’ genes *Promote*r::H2B-YFP transcriptional fusion constructs, about 2.5 to 3-kb fragment above the TSS of the *F6’H1*, *S8H* and *CYP82C4* were PCR amplified from the WT (Col-0) genomic DNA template and cloned into pENTR/D/TOPO. The resulting clones were sequence verified and used to set up an LR reaction with the gateway-compatible pGreen0229 vector, and the resulting positive clones were further confirmed by sequencing (70). Promoter reporter constructs of coumarin biosynthesis genes (*pGreen:pF6’H1*, *pGreen:pS8H*, *pGreen:pCYP82C4*) were then transformed into *Agrobacterium tumefaciens* strain GV3101. Arabidopsis WT (Col-0) and *hy5* plants were transformed using the floral dip method. T_1_ transformants were selected on ½ MS medium supplemented with glufosinate ammonium (Basta-10mg/L) and cefotaxime (100 mg/L). Single-insertion lines were identified in the T_2_ generation based on a 3:1 segregation ratio (resistant: susceptible) on selection medium (0.5 MS + Basta). Homozygous lines were screened in the T_3_ generation by confirming 100% survival on selection medium, and selected T_3_ homozygous lines were visualized under a confocal microscope.

To generate translational fusion between F6’H1, S8H and CYP82C4 and the RED FLUORESCENT PROTEIN (RFP), the full genomic sequences of these genes (i.e., including the native promoter, 5’-UTR, introns and exons but excluding the stop codon) were amplified to generate the following PCR products: *ProF6’H1:gF6’H1* (promoter length: 1787bp), *ProS8H:gS8H* (promoter length: 2232bp) and *ProCYP82C4:gCYP82C4* (promoter length: 1244bp). Each amplicon was then introduced into the entry vector pENTR4-DUAL via Gibson assembly, followed by Gateway LR recombination to transfer the constructs into the pFAST-RFP destination vector. Transformants were selected at the seed stage based on OLEOSINE1-GFP fluorescence and homozygous identified as above-described.

### YFP visualization in the transcriptional reporter lines

To visualize transcript expression *in planta,* the selected T_3_ seedlings were grown on iron-sufficient medium for 5 days, then transferred to both iron-sufficient (pH 5.7) and iron-deficient (pH 5.7 & pH 7.0) media and grown for an additional 3 days. The seedlings were stained with 10 μM propidium iodide for 1 minute. YFP fluorescence was excited using a 488-nm laser, while propidium iodide was excited with a 561-nm laser. Emission spectra were collected at 500-530 nm for YFP and 600-650 nm for propidium iodide. Imaging was conducted in Z-stack mode with a 1 μm step size, using 10× magnification and a 1.5× zoom factor on an SP8 upright confocal microscope (Leica).

### ChIP-qPCR

The ChIP assay was conducted following the protocol established by Gendrel et al. (2005) (72). Seedlings of *pHY5:HY5:YFP/hy5* and *35S:eGFP* in the WT *Ler* background were grown on 0.5 MS medium for 10 days and cross-linked with 1% formaldehyde. Chromatin was extracted and fragmented using a Qsonica 800R sonicator, applying 30 cycles of 15-second pulses at 70% amplitude, interspersed with 45-second rest intervals. HY5-DNA complexes were immunoprecipitated using protein-G magnetic beads (Dynabeads, 10004D) coupled with an anti-GFP antibody (Abcam, ab290), while rabbit IgG served as a negative control. The beads were washed, and bound complexes were eluted, then reverse cross-linked. Precipitated chromatin was analyzed by qPCR using primers listed in *SI Appendix*, Table S1, and the fold enrichment was calculated relative to the IgG control.

### Coumarin imaging

Coumarin secretion images of Arabidopsis wild type (WT) and *hy5* mutants grown in control condition (i.e., 50 μM Fe(III)NaEDTA, avFe pH 5.7) for 9 days and then transferred to media that contain a form of iron that is poorly available to plants (i.e., 100 μM FeCl_3_), at pH 5.7 (NavFe pH 5.7) or pH 7 (NavFe pH 7.0), for 3 additional days, and then observed under UV light (365 nm).

### Coumarin content estimation

Frozen root and leaf tissues were homogenized with glass beads and extracted using an 80:20 (v/v) methanol–water solution. For tissue extraction, 10–30 μL of solvent per mg fresh weight was used, maintaining a constant tissue-to-solvent ratio across all samples within each experiment. Extracts were passed through 0.45 μm filters, and equal volumes of the resulting supernatants were vacuum-dried. Dried residues were resuspended in 10 μL methanol and diluted to a final volume of 100 μL with a 90:10 (v/v) water–acetonitrile solution containing 0.1% (v/v) formic acid. Coumarin analysis was performed using high-performance liquid chromatography as previously described (73). Quantification of scopolin, scopoletin, fraxin, and esculin was achieved using six-point calibration curves generated with commercial standards (TargetMol, Boston, MA, USA; Sigma). Coumarin concentrations (nmol g^-1^ fresh weight) were calculated based on the evaporated sample volume, compound-specific response factors, and extraction solvent volume.

### HY5-coumarins spatial expression analysis

Images were obtained by linear unmixing with the emission spectra of scopolin (purple), fraxin (green), esculin (dark yellow), and propidium iodide (PI, red). The ‘Merged’ column represents the superposition of the four channels corresponding to scopolin, fraxin, esculin and PI. Seedlings were grown under control conditions (i.e., 50 μM Fe-EDTA, avFe pH 5.7) for 5 days, then transferred to control media or to media containing poorly available iron (i.e., 100 μM FeCl_3_, NavFe) at pH 5.7 or pH 7 for 6 additional days.

### Plant phenotyping assay

For phenotypic assays, the seeds were grown on half-strength iron-deficient medium, MS basal salts without iron (Murashige & Skoog basal salt mixture, Modification 33, Caisson Labs, cat. no. MSP33), supplemented with either 50 μM Fe(III)NaEDTA (ethylenediaminetetraacetic acid ferric sodium salt, Sigma-Aldrich, cat. no. E6760, CAS 15708-41-5) to provide available iron (avFe) or 50 μM FeCl (iron(III) chloride, reagent grade, Sigma-Aldrich, cat. no. 157740, CAS 7705-08-0) to simulate non-available iron (NavFe). The medium also contained 1% (w/v) sucrose and 1.2% (w/v) agar. Buffering agents were added according to pH requirements: MES for pH 5.7 and MOPS for pH 7.0, with pH adjusted using 1 M KOH. Following surface sterilization using 0.4% sodium hypochlorite followed by 70% ethanol and stratified in the dark at 4°C for 3 days, seeds were sown on pH 5.7 avFe plates and grown for 5 days. Subsequently, seedlings were transferred to media of varying pH and iron availability: pH 5.7 (avFe or NavFe) and pH 7.0 (NavFe), and grown for an additional 9 days. These plates were scanned using an Epson Perfection V600 at 1200 dpi. The root length was quantified using ImageJ 1.52a software (National Institutes of Health). For seedling growth, plants were grown under long-day conditions, 16 h light and 8 h dark, at 22°C with 50% humidity and a light intensity of 100-110 µmol cm^2^ sec^-1^.

### Chlorophyll content measurement

The chlorophyll content was measured using seedlings grown on avFe pH 5.7 and NavFe pH 5.7 and pH 7.0 for 14 days as described above. It was measured by extracting chlorophyll using 1 ml of 80% acetone from leaf tissues of three to four seedlings and incubating in the dark for 24 h using the formula: (mg/gFW) = (20.3 × A_645_ ± 8.04 × A_663_) × V/W × 10^3^ (74).

### Fe histochemical staining

Perls’ and Perls’ DAB staining (75, 76) were carried out on 5-day-old seedlings grown on 0.5 MS media at pH 5.7 and pH 7.0 to assess iron accumulation. The seedlings were vacuum-infiltrated with a solution containing 1% (v/v) HCl and 1% (w/v) K-ferrocyanide for 25 min. Seedlings were then washed with water five times, observed, and photographed using a NIKON ECLIPSE Ni U microscope. Higher Fe^3+^ levels are associated with a more intense blue colour.

For total Fe staining (Fe^3+^ and Fe^2+^), diaminobenzidine (DAB) intensification was performed after Perls’ staining. The seedlings were incubated in methanol containing 10 mM Na-azide and 0.3% (v/v) H_2_O_2_ for 1 h. The seedlings were washed with 100 mM Na phosphate buffer (pH 7.4) and incubated in the same buffer containing 0.025% (w/v) DAB and 0.005% (v/v) H_2_O_2_. The seedlings were washed with 70% ethanol to stop the reaction. The stained seedlings were then photographed using NIKON ECLIPSE Ni U microscope.

### Iron content quantification

Iron content was quantified using a spectrophotometric method based on the formation of the Fe² -BPDS complex. Whole seedlings grown under pH 5.7 and pH 7.0 with iron were harvested and separated. Approximately 30 seedlings were pooled per biological replicate, with three independent replicates analyzed. Samples were dried at 60°C for about 48 hours, and their dry weights were recorded. Dried tissues were digested in 65% (v/v) HNO at 95°C for 6 h, then treated with 30% (v/v) H O and incubated at 56°C for 2 h. An assay solution containing 1 mM bathophenanthroline disulfonate (BPDS), 0.6 M sodium acetate, and 0.48 M hydroxylamine hydrochloride was prepared. A standard curve was generated using known concentrations of FeCl . The digested samples were mixed with the assay solution to form a pink-colored Fe² -BPDS complex, and absorbance was measured at 535 nm. Iron concentrations were calculated from the standard curve and normalized to the sample dry weight (77, 78).

### Co-localization studies

Translational reporter line of HY5 (*pHY5:HY5:YFP*) was crossed with coumarin translational reporter lines (*pF6’H1:F6’H1:RFP*, *pS8H:S8H:RFP*, *pCYP82C4:CYP82C4:RFP* and *pBGLU42:BGLU42:RFP*). The F_1_ seeds obtained were grown in Fe-sufficient conditions (+Fe, 50µM Fe(III)NaEDTA, pH 5.7) for 4 days and then transferred to +Fe and Fe-deficient conditions (-Fe, pH 5.7 and -Fe, pH 7.0) for 3 days. The fluorescent images were obtained using a confocal laser scanning microscope SP8 (Leica). YFP and RFP were excited at 488 and 561 nm and the signals were detected between 500-550 nm and 610-660 nm, respectively. YFP and mCitrine share similar excitation and emission spectra; therefore, the fluorescent images of *pHY5:HY5:YFP* x *pPDR9:PDR9:mCitrine* seeds were obtained by exciting YFP and differentiated based on the localization of the two proteins, HY5 in the nucleus and PDR9 in the plasma membrane.

### Fraxetin Supplementation assay

The plants were grown in media containing Fe that is poorly available to plants (100 µM FeCl_3_) at pH 7.0 for 4 days, then transferred to media containing 100 µM FeCl_3_ at pH 7.0, supplemented with or without 50 µM Fraxetin, for 3 days. Sterile DMSO was added to the media not supplemented with fraxetin. Fresh stock solution of fraxetin was prepared in DMSO and filter-sterilized before use. Primary root length and chlorophyll content were measured as already described.

### Statistical analysis

The graphs were plotted using GraphPad PRISM 11. Significance was calculated using One-way ANOVA followed by a post hoc Tukey HSD test or Two-way ANOVA followed by a Tukey’s multiple comparison test. Different letters (a, b, c) were used to indicate significant differences. Error bars represented the ± SEM. A p-value ≤ 0.05 was considered statistically significant.

## ACKNOWLEDGMENTS AND FUNDING SOURCES

We are thankful to all the members of SBS and CD labs. SBS acknowledges intramural funding support from Indian Institute of Science Education and Research (IISER) Mohali. SBS acknowledges the support through grant no. BT/PR51324/AGIII/103/1480/2023 from the Department of Biotechnology (DBT). SBS also acknowledges the Science and Engineering Research Board (SERB) for research funding (CRG/223 2022/003773). The SBS laboratory is also supported by the Indo French Centre for the Promotion of Advanced Research (IFCPAR/ CEFIPRA) under project 68T06 1. VK is supported by a PhD fellowship from the Indian Institute of Science Education and Research (IISER) Mohali. CD thank Dr. Carine Alcon and Matthieu Déjean for technical assistance and expertise for microscope observations, and the imaging facility MRI, member of the national infrastructure France-BioImaging supported by the French National Research Agency (ANR-10-INBS-04, ‘Investments for the future’). CD also acknowledges the French National Research Agency (DYNAFER project, ANR-22-CE20-0006) and the National Research Institute for Agriculture, Food and the Environment (INRAE, BAP Department, TRACE project). MK was supported by a PhD fellowship from the European teaming program Biopolis (HOMEOFER project), ML by the China Scholarship Council and SW by a Marie Skłodowska-Curie Individual Fellowship in Horizon 2020 from the European Council (PLANTSEE project, MSCA-IF-2020, 101024030).

## COMPETING INTERESTS

The authors declare no competing interests.

**Figure S1. *HY5* positively regulates expression of coumarin secretion genes.**

(*A-B*) RT-qPCR of coumarin secretion genes *PDR9* and *BGLU42* in WT and *hy5* seedlings. Seedlings were grown on +Fe pH 5.7 media for 6 days and then transferred to +Fe pH 5.7, - Fe pH 5.7, and -Fe pH 7.0 for 3 days. Transcript levels were normalized to *PP2AA3*. Data are mean ± SEM of three biological replicates (two technical replicates each); each biological replicate is pooled RNA from ∼100 roots. Different letters indicate significant differences (one-way ANOVA, Tukey’s post hoc test, p < 0.05). (*C*) Schematic of the *BGLU42* promoter region; the line below the box indicates the region analyzed by ChIP-qPCR, with distance upstream of the start codon shown in brackets. (*D*) ChIP-qPCR enrichment of HY5 at the BGLU42 promoter region in (C), using *pHY5::HY5:YFP/hy5* (Ler) and *35S::eGFP* (Ler) seedlings grown on +Fe for 10 days. n = 2 biological replicates per genotype. Data are mean ± SEM. Different letters indicate significant differences (one-way ANOVA, Tukey’s post hoc test, p < 0.05).

**Figure S2. *HY5* positively regulates the expression of and directly binds to the promoters of the coumarin regulatory genes *MYB10* and *MYB72*.**

(*A-B*) RT-qPCR of *MYB10* and *MYB72* in WT and *hy5* seedlings. Seedlings were grown on +Fe pH 5.7 media for 6 days and then transferred to +Fe pH 5.7, -Fe pH 5.7, and -Fe pH 7.0 for 3 days. Transcript levels were normalized to *PP2AA3*. Data are mean ± SEM of three biological replicates (two technical replicates each); each biological replicate is pooled RNA from ∼100 roots. Different letters indicate significant differences (one-way ANOVA, Tukey’s post hoc test, p < 0.05). (*C-D*) Schematics of the *MYB10* and *MYB72* promoter regions; lines below each box indicate the region analyzed by ChIP-qPCR, with distances upstream of the start codon shown in brackets. (*E-I*) ChIP-qPCR enrichment of HY5 at the *MYB10* and *MYB72* promoter regions in (C-D), using *pHY5::HY5:YFP/hy5*and *35S::eGFP* seedlings grown on +Fe for 10 days. n = 2 biological replicates per genotype. Data are mean ± SEM. Different letters indicate significant differences (one-way ANOVA, Tukey’s post hoc test, p < 0.05).

**Figure S3. HY5 and coumarin pathway genes act in a common genetic pathway to sustain chlorophyll content under high pH**

(*A*) Scatter plot showing the mean of the total chlorophyll content estimated in WT*, hy5, f6’h1, f6’h1 hy5, s8h*, *s8h hy5, cyp82c4*, *cyp82c4 hy5, pdr9,* and *pdr9 hy5*. Five days old seedlings grown at pH 5.7 on avFe were transferred to avFe at pH 5.7 and NavFe media at pH 5.7 and pH 7.0, and grown for an additional 9 days before chlorophyll content estimation. Each replicate consists of 3 shoots pooled for estimation. n∼6-9 biological replicates. Means with different letters are significantly different according to two-way ANOVA followed by a post hoc Tukey test, p < 0.05, using GraphPad PRISM 11. Error bars represent ±SEM.

**Figure S4. *HY5* genetically interacts with the coumarin regulatory gene *MYB72* involved in iron acquisition at high pH**

(*A*) Phenotype (scale bar=1cm), (*B*) boxplots of primary root length (n depicted in the x-axis beneath corresponding samples), and (*C*) chlorophyll content of WT*, hy5, myb72,* and *myb72 hy5* seedlings. Five days old seedlings grown at pH 5.7 on avFe were transferred to avFe at pH 5.7 and NavFe media at both pH 5.7 and pH 7.0 and grown further for nine days before data acquisition. Means with different letters are significantly different according to two-way ANOVA followed by a post hoc Tukey test, p < 0.05, using GraphPad PRISM 11. Error bars represent ±SEM.

**Figure S5. Coumarin biosynthesis and secretion genes and *HY5* are involved in the same pathway for iron acquisition**

(*A*) PERLS’ and (*B*) PERLS’-DAB stained maturation zone images of five days old WT*, hy5, f6’h1, f6’h1 hy5, s8h*, *s8h hy5, cyp82c4*, *cyp82c4 hy5, pdr9* and *pdr9 hy5* seedlings grown on iron sufficient medium (+Fe) at both pH 5.7 and pH 7.0. A representative image from three independent experiments, with ∼10 seedlings in each condition, is shown. Scale bar = 100 μm. (*C*) Iron content was quantified in whole seedlings grown on +Fe media for 15 days. The data represent the average of three biological replicates, each with three technical replicates. Each biological replicate consisted of ∼30 whole seedings. Error bars represent ±SEM. DW - dry weight. Different letters (a, b, c, d) indicate significant differences, determined by two-way ANOVA followed by a Tukey’s multiple comparison test (P≤0.05) using GraphPad PRISM 11.

**Figure S6. Altered iron accumulation WT, *hy5*, *myb72* and *myb72 hy5* at high pH**

(*A*) Perls’ and (*B*) Perls’-DAB stained maturation zone images of five-day-old WT, *hy5*, *myb72,* and *myb72 hy5* seedlings grown on iron sufficient medium (+Fe) at both pH 5.7 and pH 7.0. A representative image from three independent experiments, with ∼10 seedlings in each condition, is shown. Scale bar = 100 μm. (*C*) Iron content in whole seedlings grown on +Fe for 15 days. Data are mean ± SEM of three biological replicates (three technical replicates each); each biological replicate is ∼30 pooled seedlings. DW, dry weight. Different letters indicate significant differences (two-way ANOVA, Tukey’s post hoc test, p ≤ 0.05).

**Figure S7. HY5 colocalizes with BGLU42.**

*A)* The seedlings of HY5 translational YFP fusion line crossed with BGLU42 translational RFP fusion lines were grown on +Fe medium for four days and transferred to -Fe medium at pH 5.7 and pH 7.0 for another 3 days. Yellow: YFP, magenta: RFP. Confocal imaging was performed in Z-stack mode with a step size of 1 µm. Scale bar= 10 µm.

**Figure S8. Fraxetin restores chlorophyll content under high pH mediated iron deficiency**

(*A*) Total chlorophyll content of WT*, hy5, f6’h1, f6’h1 hy5, s8h, s8h hy5, cyp82c4, cyp82c4 hy5, pdr9* and *pdr9 hy5* grown on NavFe media at pH 7.0. for four days and transferred to NavFe at pH 7.0 media with fraxetin or solvent control (DMSO) and grown further for three days. Means with different letters are significantly different according to one-way ANOVA followed by a post hoc Tukey test, p < 0.05, using GraphPad PRISM 11. Error bars represent ±SEM.

