## Supplementary figures and images for "HY5 orchestrates the transcriptional network controlling coumarin-mediated iron acquisition under elevated pH conditions"

### Supplementary figures

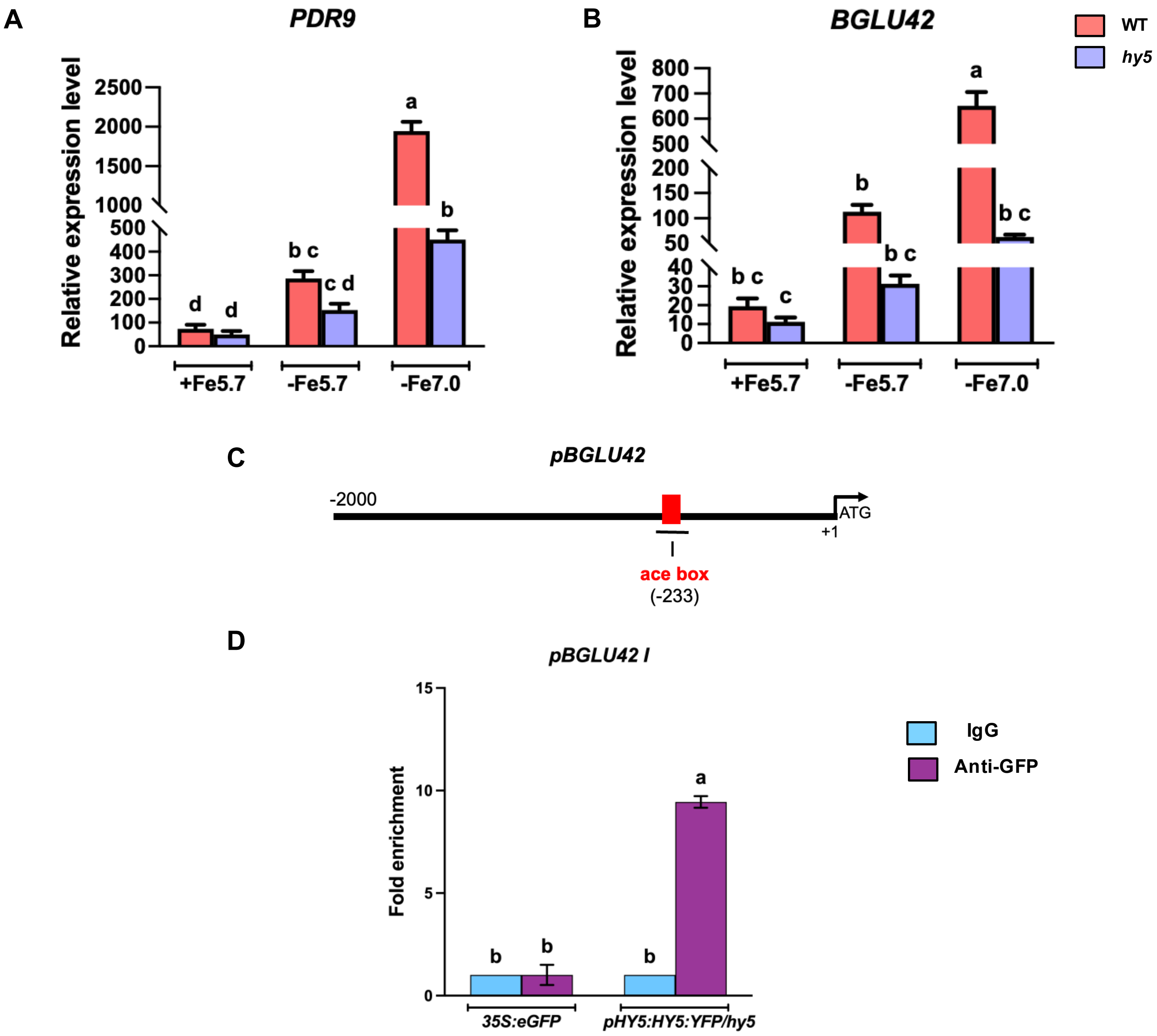

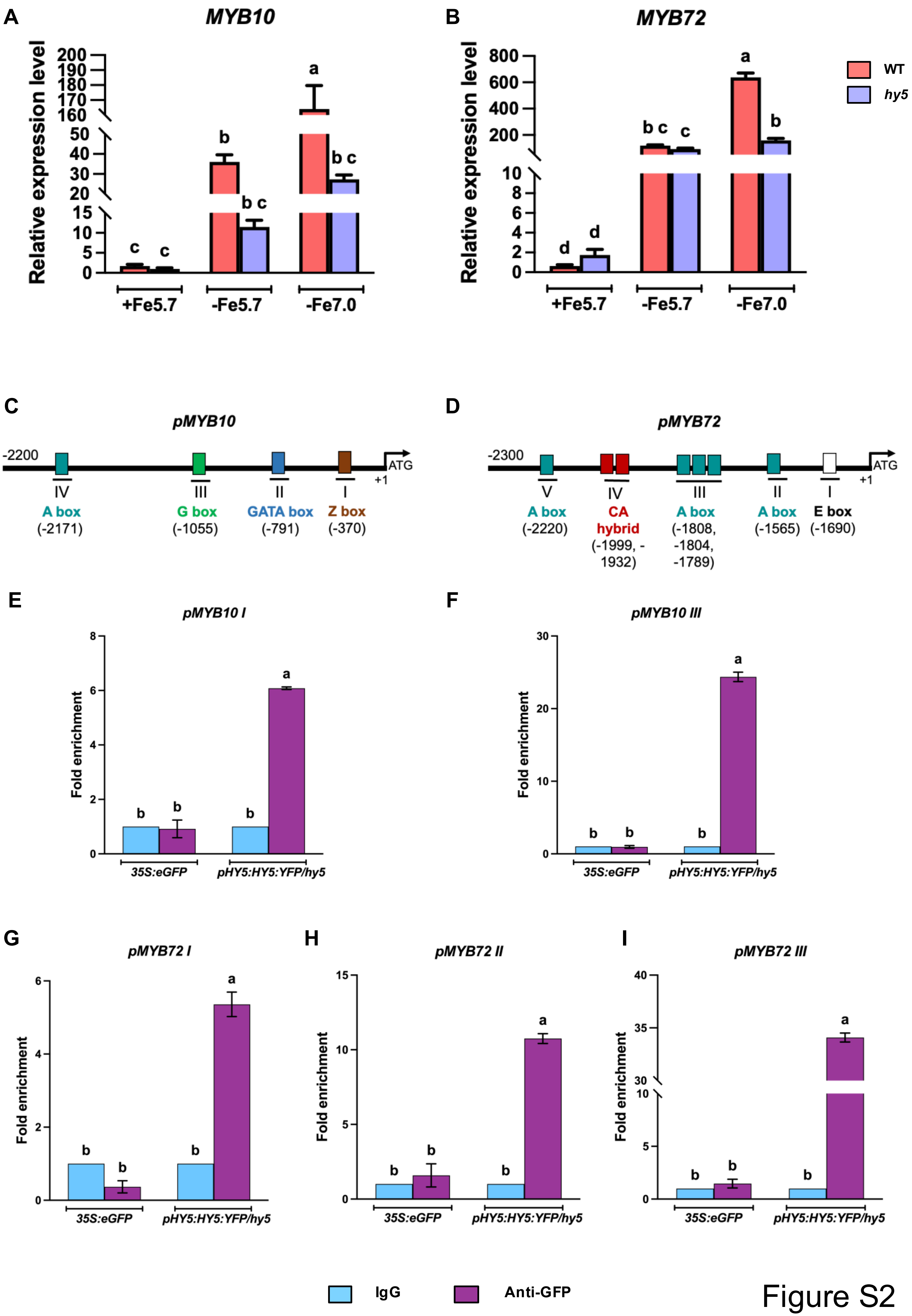

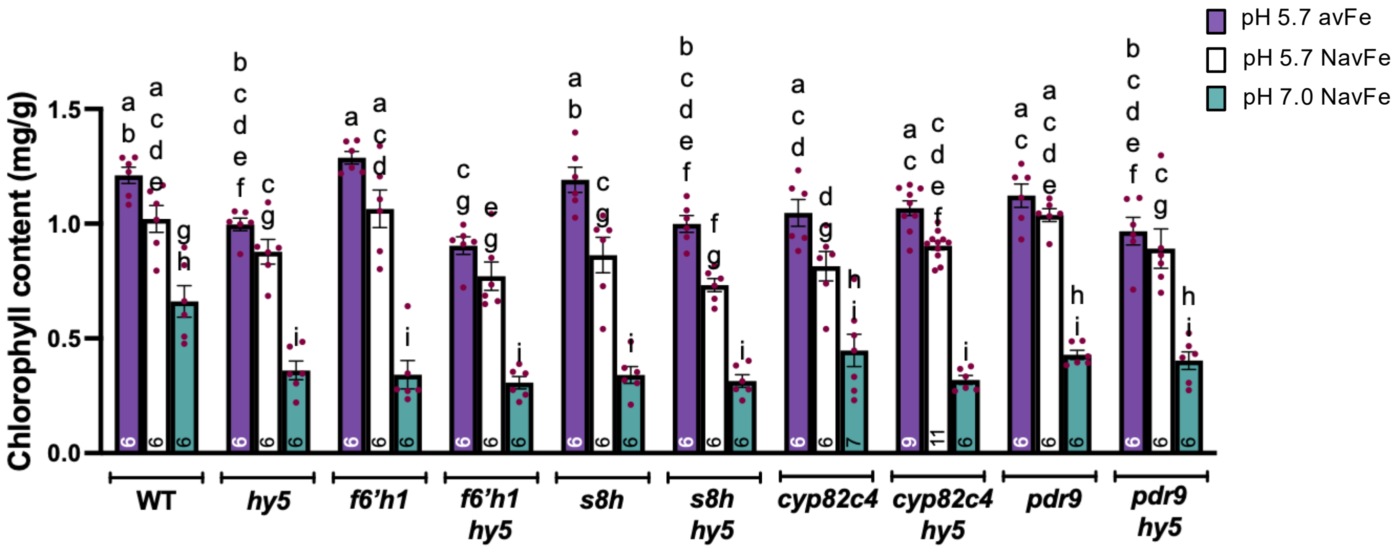

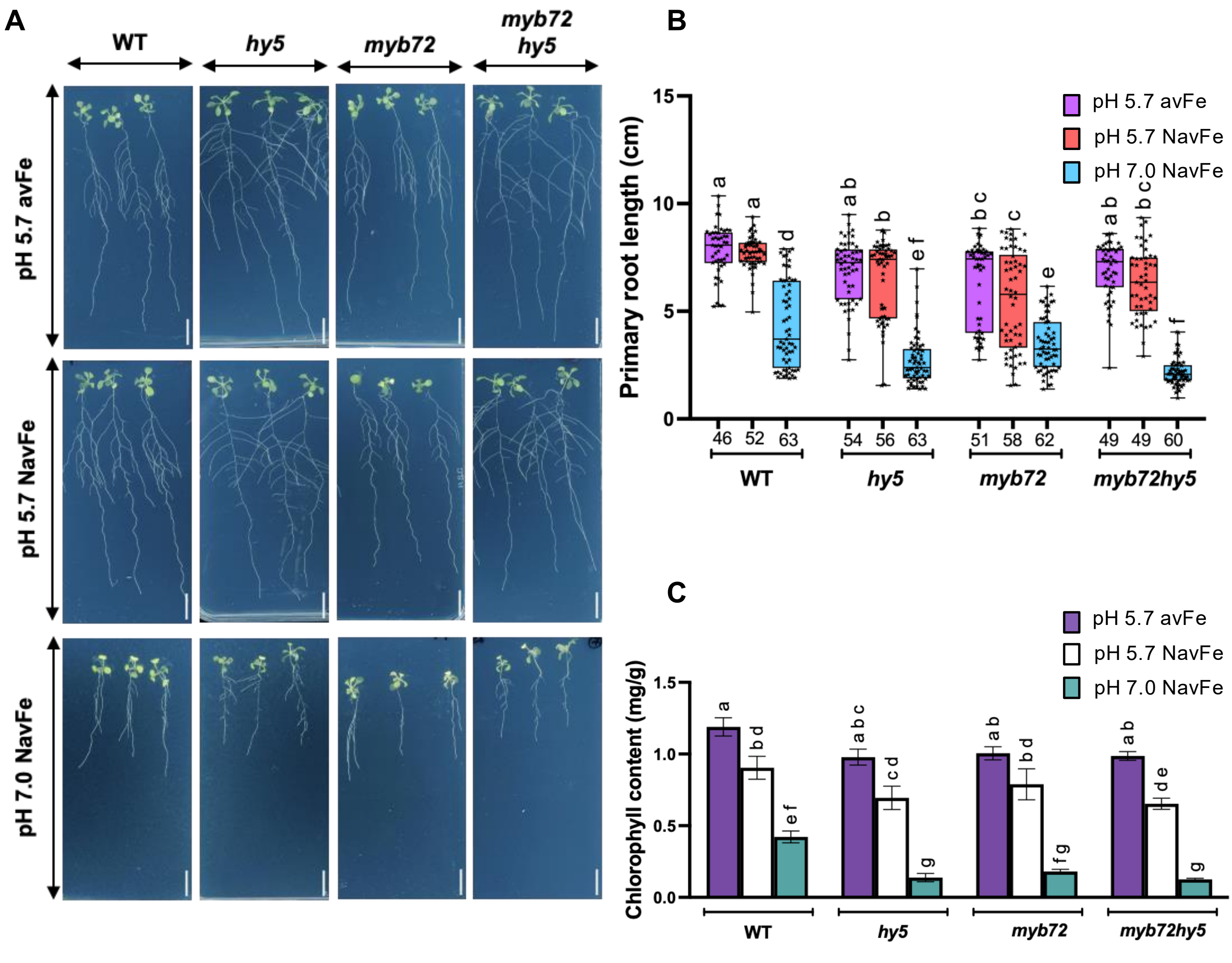

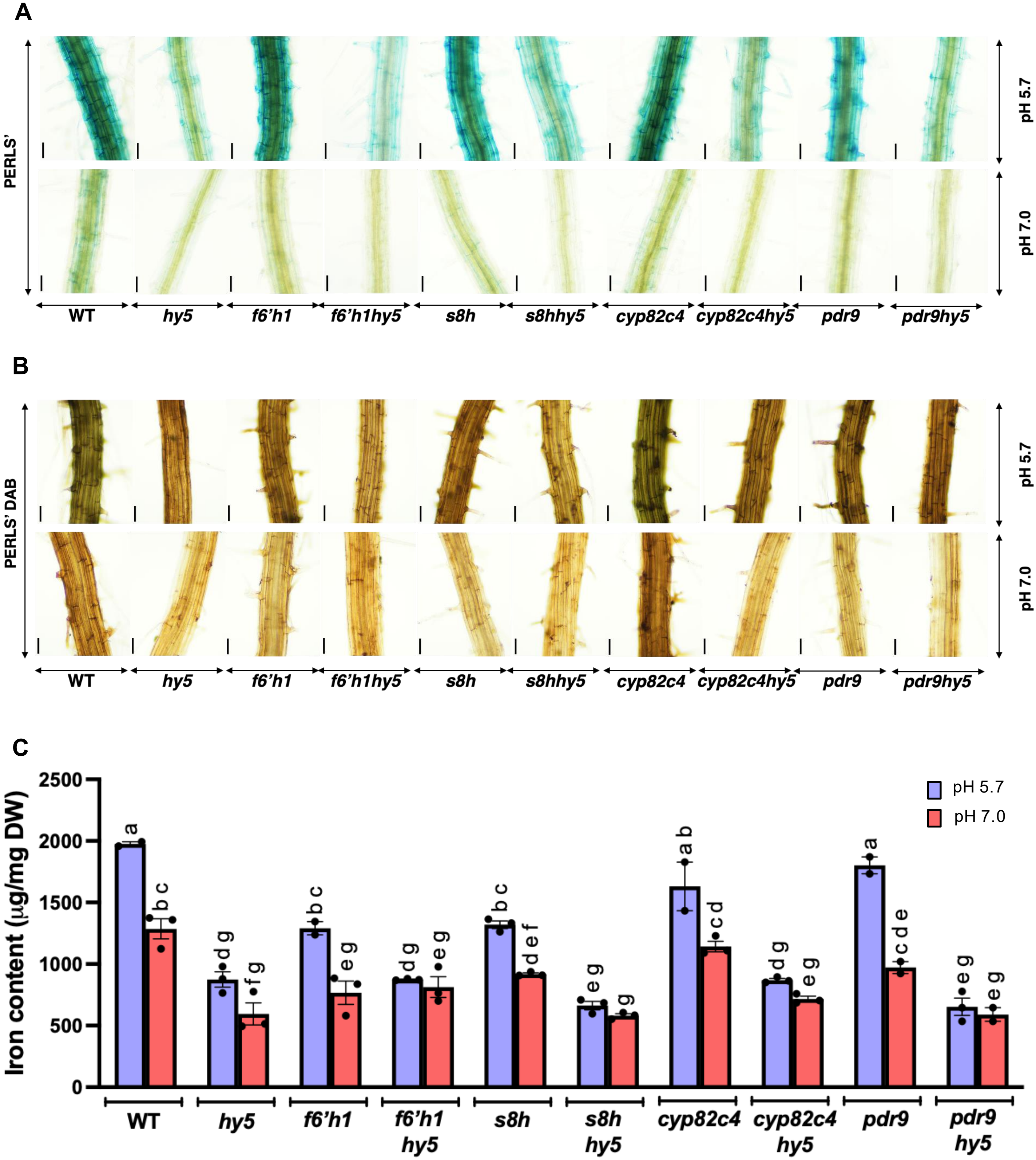

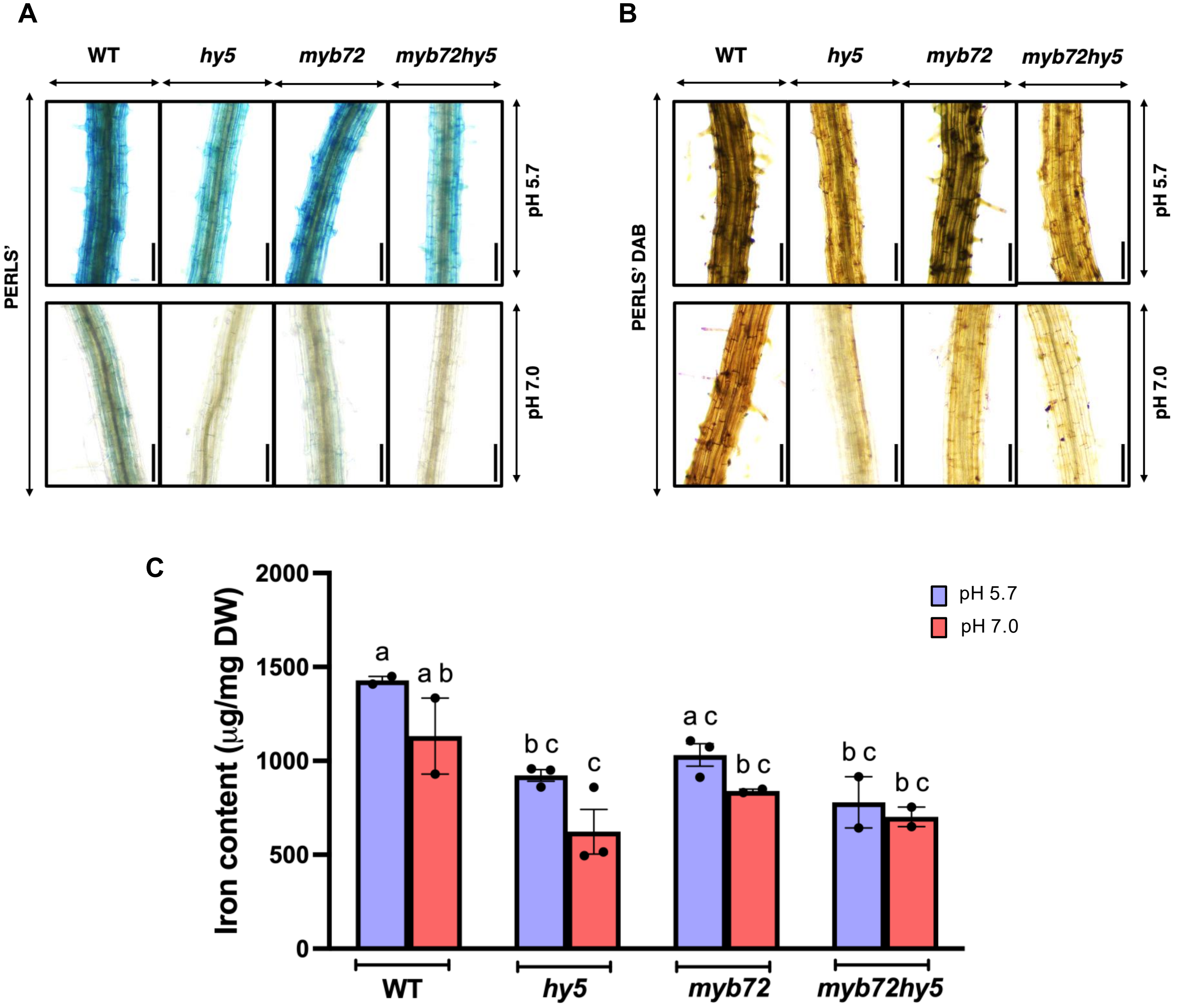

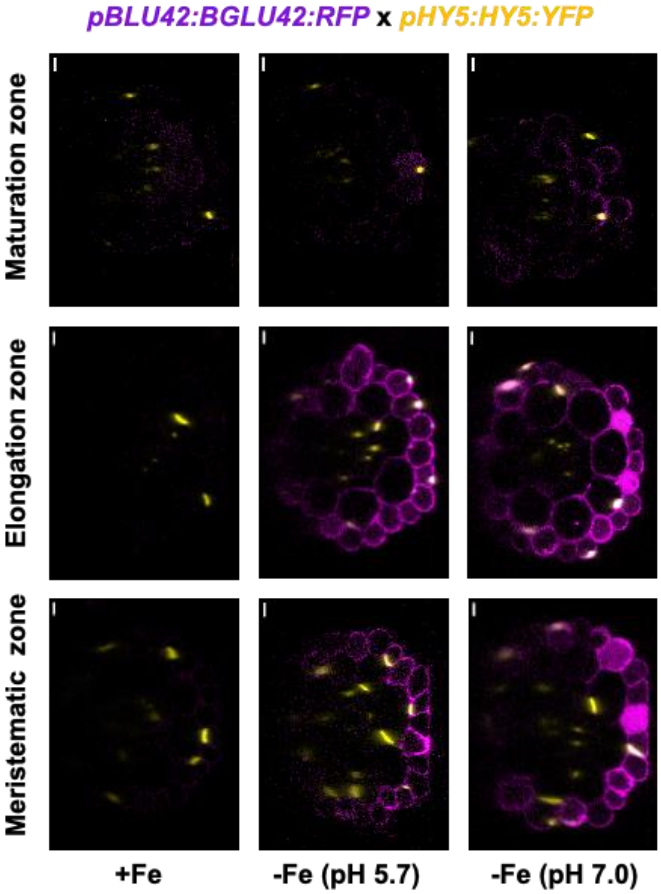

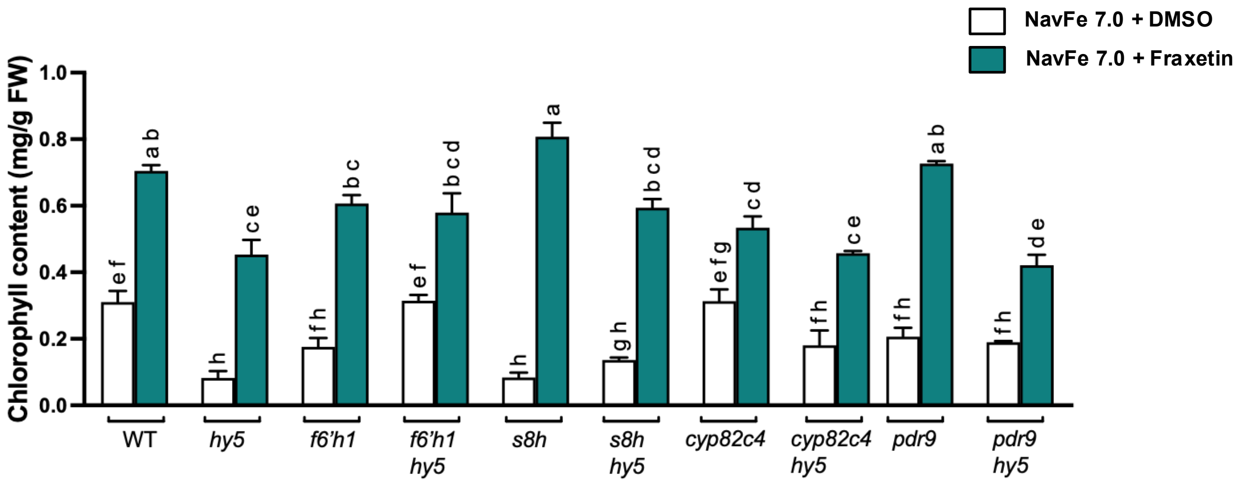
